# Thermosensitivity of cellular translation restricts the growth of fission yeast at high temperatures

**DOI:** 10.1101/2025.08.07.669215

**Authors:** Yutaka Akikusa, Atsuya Yamaguchi, Yutaka Funahashi, Fontip Mahayot, Taisuke Naka, Ami Matsuo, Yukiko Nakase, Shingo Izawa, Kazuhiro Shiozaki, Yuichi Morozumi

## Abstract

Living organisms have thermal limits above which they are unable to operate and survive. Our previous genetic screen identified proteins that impede the high-temperature growth of fission yeast, including the RNA-binding protein Dri1 and a fission yeast-specific protein termed Rhs1. Here, we show that Dri1 and Rhs1 form a complex and physically interact with the Ccr4-Not complex, a master regulator of mRNA metabolism. Gene expression analysis revealed that the Dri1-Rhs1 and Ccr4-Not complexes negatively regulate a set of genes implicated in ribosome biogenesis (Ribi genes). Loss of the Dri1-Rhs1 complex results in the augmented expression of Ribi genes, thereby suppressing the accumulation of 80S monosomes and the growth inhibition under high-temperature conditions. The thermosensitivity of the translational processes may be a determinant of the upper limit of the growth temperature in fission yeast.

## Introduction

Elevated temperatures are one of the major stress conditions that can impair the growth and development of living organisms by causing protein denaturing and heat damage to other cellular macromolecules. To counteract and survive such stressful conditions, cells activate the heat shock response (HSR). This evolutionarily conserved mechanism reprograms gene expression and induces the synthesis of heat shock proteins (HSPs). Most HSPs are known to function as molecular chaperones, which maintain cellular proteostasis by refolding denatured proteins, preventing their aggregation, and sorting unfolded proteins for degradation^1,2^. Hsp104, an HSP that dissolves protein aggregates and facilitates protein refolding, is necessary for cell survival in response to acute heat stress in *Saccharomyces cerevisiae* and *Candida albicans*^3–5^.

In contrast, budding yeast *UBI4*, a polyubiquitin-encoding gene inducible under high-temperature conditions, is dispensable for the acute heat stress response but is essential for adaptation to chronic, sublethal high temperatures^6,7^. It was also recently reported that tolerance to short-term and long-term heat stress in *Arabidopsis thaliana* requires distinct sets of factors^8–10^. These observations strongly suggest that cells employ different adaptive mechanisms depending on the intensity and duration of heat stress. While pro-survival mechanisms for acute heat shock, including the HSR, have been extensively studied, the molecular mechanisms that contribute to thermotolerance to chronic heat stress remain poorly understood.

The optimal growth temperature of the fission yeast *Schizosaccharomyces pombe* is around 30°C, and its growth is severely impaired at 38°C and above. We previously found that *S. pombe* cells proliferate even at 39°C when the activity of TOR complex 1 (TORC1), a kinase complex conserved among diverse eukaryotes, is suppressed^11^. Consistently, the null mutants of Sck1 and Mks1, both of which are the TORC1 substrates, exhibit significant cell growth at 39°C. These results indicate that the TORC1 signaling pathway restricts the high-temperature growth of fission yeast cells.

We also conducted genetic screens in *S. pombe* for additional negative regulators of cell proliferation at high temperatures and identified Dri1 and Rhs1^11^. Dri1 is an RNA-binding protein implicated in heterochromatin assembly and kinesin loading to the mitotic spindle^12,13^. On the other hand, Rhs1 is a fission yeast-specific protein whose molecular function remains unknown. The high-temperature growth of a fission yeast strain lacking both the *dri1* and *rhs1* genes is comparable to that of the respective single mutants. Moreover, a physical interaction between Dri1 and Rhs1 was observed, suggesting that Dri1 and Rhs1 function as a complex in the negative regulation of high-temperature cell growth^11^.

The Ccr4-Not complex is a eukaryotic protein complex that serves as a master regulator of mRNA metabolism, from transcription to degradation, thereby modulating gene expression^14,15^. The fission yeast Ccr4-Not complex contains seven highly conserved subunits, including the scaffold protein Not1 as well as the Ccr4 and Caf1 deadenylases that degrade the poly(A) tail of mRNA. The other subunits are Mot2, Not2, Not3, and Rcd1. Mot2 is a RING finger E3 ubiquitin ligase, and Not2 and Not3 contain the NOT box motif. Little is known about the function of the Rcd1 subunit within the complex. The null mutation of either *caf1*, *ccr4,* or *mot2* causes growth defects in fission yeast^16–18^, indicating that the function of Ccr4-Not is important for normal cell growth. On the other hand, the involvement of this complex in the cellular adaptation to temperature changes has never been examined.

In this study, we have demonstrated that the complex formation between Dri1 and Rhs1 is essential for their function, and its disassembly results in cell proliferation even at growth-inhibitory, high temperatures. A search for factors physically interacting with the Dri1-Rhs1 complex has identified the Ccr4-Not complex. Indeed, the null mutation of some genes encoding Ccr4-Not subunits confers cellular heat resistance, suggesting that the interacting two complexes, Dri1-Rhs1 and Ccr4-Not, restrict cell proliferation at high temperatures. Interestingly, genes implicated in ribosome biogenesis (Ribi genes) are up-regulated in strains lacking the Dri1-Rhs1 complex or the Ccr4-Not subunit Not3. The augmented expression of Ribi genes in those mutant strains ameliorates the 80S monosome accumulation and growth defects under high-temperature conditions. These results have shed light on protein translation as a thermosensitive process that affects the upper-temperature limit for cell growth.

## Results

### Residues 31-50 of Rhs1 mediate its interaction with Dri1

We previously identified fission yeast Dri1 and Rhs1 as suppressors of cell growth at high temperatures; the *dri1* and *rhs1* null mutants can proliferate at 39°C, where wild-type cells arrest their growth and lose viability^11^. Although physical interaction between Dri1 and Rhs1 was detected, it remains to be determined whether they function as a protein complex in restricting high-temperature growth. We, therefore, set out to identify the amino acid sequence within Rhs1 required for the interaction with Dri1 by yeast two-hybrid assays. Although autoactivation was observed when Rhs1 was used as bait in the assay, the Dri1-Rhs1 interaction was successfully detected by utilizing Dri1 and Rhs1 as bait and prey, respectively (Fig. S1). Dri1 interacted with the C-terminally truncated fragment of Rhs1(1-240) but not the N-terminally truncated one (121-368) (Fig. 1A). Consistently, the N-terminal 120 amino residues of Rhs1(1-120) were sufficient for the interaction with Dri1. Moreover, these observations were confirmed by immunoprecipitation experiments with fission yeast strains expressing a series of truncated Rhs1; Dri1 was co-purified with the FLAG-tagged Rhs1 fragments containing residues 1-120 (Fig. 1B). Thus, the N-terminal 120 residues of Rhs1 include the Dri1-binding region.

**Fig. 1.**
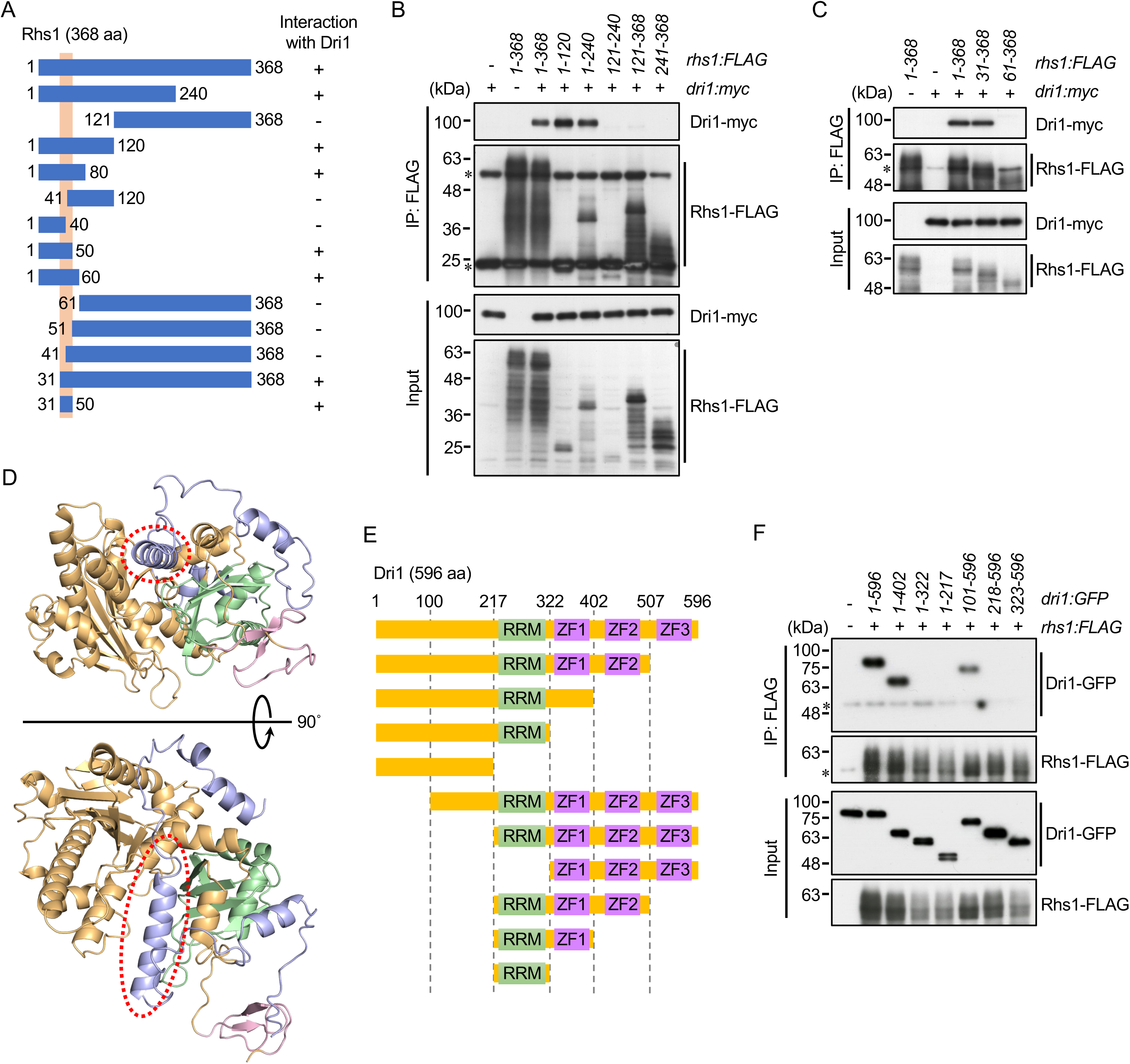
The residues 31-50 of Rhs1 are indispensable for the interaction with Rhs1. (A) Summary of the interaction between Dri1 and various Rhs1 fragments in yeast two-hybrid assays. Interaction was monitored by *ADE2* and *HIS3* reporter gene expression in the budding yeast Y2HGold strain as shown in Fig. S1A. +, interaction; -, no interaction. An orange box indicates the Rhs1 region required for the Dri1-Rhs1 interaction. (B and D) Physical interaction between Dri1 and the various truncated Rhs1 fragments. *dri1:myc* cells expressing various Rhs1 fragments were cultured in YES medium at 30°C and their cell lysate was subjected to immunoprecipitation using the anti-FLAG antibodies (IP: FLAG), and co-purified Dri1-myc was analyzed by immunoblotting. (C and F) The growth of the strains expressing the various Rhs1 fragments at high temperatures. The indicated *rhs1* mutant strains were grown in YES medium at 30°C, and their serial dilutions were spotted onto YES agar plates for growth assay at 30°C and 39°C. (E) Predicted structure of Rhs1 (light blue) binding to Dri1 (light orange) using Alphafold 3. The RRM and RanBP2-type zinc finger of Dri1 are shown in pale green and light pink, respectively. Disordered regions of Dri1 (364-596) and Rhs1 (96-368), which do not form any interactions, are hidden in the structure. A dashed oval (red) indicates the region of Rhs1 (31-50) required for the interaction with Dri1 in yeast two-hybrid assay and immunoprecipitation. (G) A schematic diagram of the Dri1 fragments. RRM, RNA recognition motif; ZF1-3, RanBP2-type zinc finger. (H) Physical interaction between Rhs1 and the various truncated Dri1 fragments. *rhs1:FLAG* cells expressing various Dri1 fragments were cultured in YES medium at 30°C and their cell lysate was subjected to immunoprecipitation as in (B). (I) High-temperature growth of the strains expressing the various Dri1 fragments. The indicated *dri1* mutant strains were grown in YES medium at 30°C, and their serial dilutions were spotted onto YES agar plates for growth assay at 30°C and 39°C.

We repeated the yeast two-hybrid assay to further narrow down the Dri1-binding region. The Dri1 interaction was detected with the Rhs1 N-terminus of 50 residues or longer (Fig. 1A). In addition, while the Rhs1 fragment lacking the N-terminal 30 residues (31-368) interacted with Dri1, further truncation of N-terminal 40 residues or more abrogated the association in the yeast two-hybrid and immunoprecipitation assays (Fig. 1C). Indeed, the Rhs1 fragment of only residues 31-50 was sufficient for the interaction with Dri1 (Fig. 1A). Notably, in a structure prediction of the Dri1-Rhs1 complex by AlphaFold 3 (Fig. 1D; Abramson et al., 2024), this region of Rhs1 forms an α-helix and is inserted into a groove formed by the N-terminal region and RNA recognition motif (RRM) of Dri1 (see below). Together, these results strongly suggest that residues 31-50 of Rhs1 contain the sequence to bind Dri1.

Dri1 harbors a single RRM and three repeats of the RanBP2-type zinc finger domain (ZF1-3), both of which are known as RNA-binding domains (Fig. 1E). As residues 31-50 of Rhs1 are predicted to bind the groove formed between the N-terminal region and RRM of Dri1 (Fig. 1D), we next tested this interaction model by immunoprecipitation using strains expressing a series of truncated Dri1 proteins (Fig. 1F). Like full-length Dri1, the Dri1 fragment lacking the two zinc fingers from the C-terminus (1-402) was co-purified with Rhs1 (Fig. 1F). The Rhs1 interaction was partially compromised with Dri1(101-596), which lacks the N-terminal 100 residues that are not part of the groove in the predicted structure (Fig. 1D). On the other hand, the N-terminal fragment without RRM (1-217) and Dri1(218-596) lacking the entire N-terminal region failed to bind Rhs1, confirming the contribution of both the N-terminal region and RRM to the Dri1-Rhs1 interaction. However, the Dri1(1-322) fragment, consisting only of the N-terminal domain and RRM, failed to bind to Rhs1 (Fig. 1F); such a minimal fragment might not be properly folded to form the Rhs1-binding groove.

### Formation of the Dri1-Rhs1 complex is essential for restricting high-temperature growth

As shown above, the Dri1-binding region within Rhs1 was successfully narrowed down to its residues 31-50 (Fig. 1A). To identify the amino acid residues essential for Dri1 binding, a series of Rhs1 mutants were constructed by replacing two consecutive residues in this Rhs1 sequence with alanine (Fig. 2A). Among the Rhs1 mutant proteins constructed, only Rhs1-AF33.34AA and -QF37.38AA failed to interact with Dri1 in the yeast two-hybrid assay (Fig. 2A). Consistently, the amount of co-purified Dri1 was significantly decreased when FLAG-tagged Rhs1-FQF34.37.38AAA, -FQ34.37AA, and -FF34.38AA were immunoprecipitated (Fig. 2B). These results were further assessed by constructing the alanine mutants of the individual residues. Single substitutions at Phe-34 and Gln-37 in Rhs1 drastically abrogated the copurification of Dri1, while that of Phe-38 only partially compromised the Dri1-Rhs1 interaction (Fig. 2B). These results indicate that the Phe-34 and Gln-37 of Rhs1 are the critical residues for the interaction with Dri1.

**Fig. 2.**
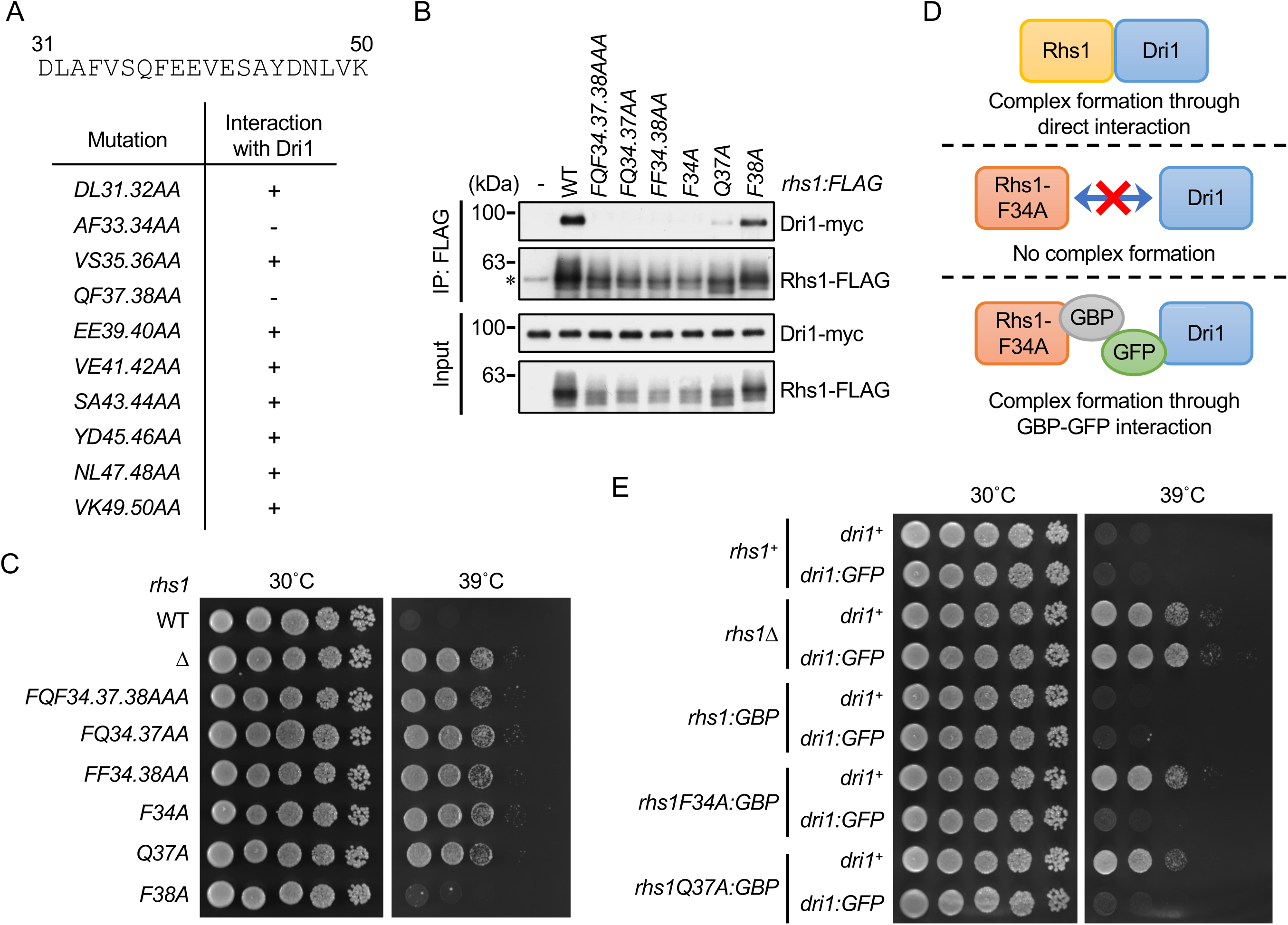
Formation of the Dri-Rhs1 complex is indispensable for its function in the negative regulation of high-temperature growth. (A) Interaction between Dri1 and the Rhs1 mutants carrying different alanine substitutions in the 31-50 residues of Rhs1 was examined in yeast two-hybrid assays as in Fig. 1A. The amino acid sequence of Rhs1(31-50) is shown at the top. (B) Phe-34, and Gln-37 of Rhs1 are critical for its interaction with Dri1. *dri1:myc* cells expressing the indicated alanine mutants of Rhs1 were cultured in YES medium at 30°C and their cell lysate was subjected to immunoprecipitation as in Fig. 1B. (C) The F34A and Q37A substitutions in Rhs1 confer cellular heat resistance on fission yeast. The indicated *rhs1* mutant strains were grown in YES medium at 30°C, and their growth was tested at indicated temperatures by spotting serial culture dilutions on YES agar plates. (D) A schematic representation of the Dri1-Rhs1 complex formation mediated by the GFP-GBP interaction. (E) Dri1 and Rhs1 function as a complex for suppressing cell growth at high temperatures. The indicated strains were grown in YES medium at 30°C, and their growth was tested as in (C).

The functional significance of the Dri1-Rhs1 complex formation was evaluated by the high-temperature growth assay of strains expressing the alanine-substitution mutants of Rhs1 described above. The *rhs1* mutants carrying *F34A* and/or *Q37A*, both of which abrogate the Dri1-Rhs1 interaction (Fig. 2B), exhibited heat-resistant growth similar to that of *rhs1Δ* (Fig. 2C). On the other hand, a strain expressing Rhs1-F38A, which still interacts with Dri1 (Fig. 2B), failed to grow at 39°C, like the wild type. Thus, the loss of the Dri1-Rhs1 association results in heat resistance, as observed with the *dri1Δ* and *rhs1Δ* mutants. Moreover, we found that the *rhs1F34A* and *rhs1Q37A* defects were suppressed by artificially tethering the Rhs1 mutant proteins to Dri1 through the GFP-GFP binding protein (GBP) targeting system^20^ (Fig. 2D). When Rhs1 with *F34A* or *Q37A* mutations was expressed as GBP-fusions together with GFP-fused Dri1, the cells exhibited the wild-type phenotype and were sensitive to high temperatures (Fig. 2E). Together, these results strongly suggest that the association between Dri1 and Rhs1 is indispensable for their functionality in restricting high-temperature cell growth.

### Dri1, but not Rhs1, forms cytoplasmic aggregates upon high-temperature stress

We examined the cellular localization of Dri1 and Rhs1 by fluorescence microscopy of strains expressing these proteins tagged with GFP from their chromosomal loci. Dri1 was found to diffuse throughout the cytoplasm (Fig. 3A). The protein appears largely excluded from the nucleus, except for a single, dot-like signal within the nucleus. Interestingly, the *rhs1Δ* mutation resulted in the distribution of Dri1 throughout the nucleus and cytoplasm, and brighter, multiple Dri1 puncta were detected in the nucleus (Fig. 3A).

**Fig. 3.**
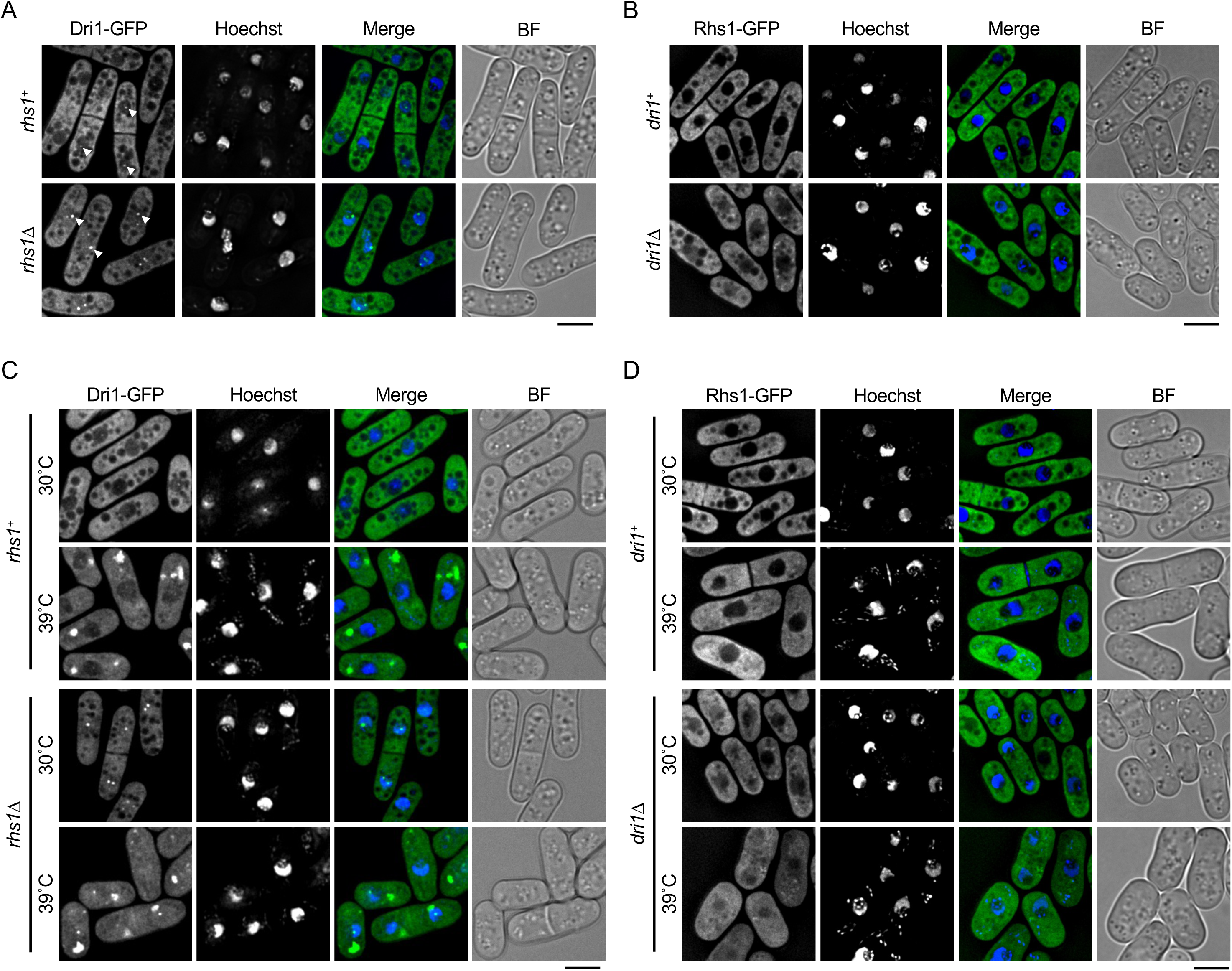
Dri1 and Rhs1 regulate their cellular localization by one another. (A and B) Indicated fission yeast strains expressing GFP-tagged Dri1 or Rhs1 protein from their chromosomal locus were grown in EMM at 30 C°, and its localization was analyzed by fluorescence microscopy. The nucleus was visualized by Hoechst staining. Z-axial images were collected and mid-section images after deconvolution are shown. BF, bright-field image. Bars, 5 µm. (C and D) Strains used in (A) and (B) were grown in EMM at 30 C° and shifted to 39°C. After 4-hour incubation at 39°C, cells were subjected to fluorescence microscopy as in (A) and (B).

As shown in Figure 3B, Rhs1-GFP was diffused in the cytoplasm but excluded from the nucleus; unlike Dri1-GFP, no nuclear puncta were observed. In the *dri1Δ* background, some diffused Rhs1 signals were detectable in the nucleus. One possible interpretation for these observations is that Dri1 and Rhs1 are excluded from the nucleus once they are assembled into a complex; however, a fraction of Dri1 forms nuclear foci independently of Rhs1.

The cellular localization of Dri1 and Rhs1 was also examined under high-temperature conditions. After a temperature shift to 39°C, Dri1 formed cytoplasmic aggregates both in the *rhs1*^+^ and *rhs1Δ* backgrounds (Fig. 3C). A recent study^13^ reported that Dri1 is incorporated into Protein Aggregate Centers (PACs), cytoplasmic protein aggregates believed to sequester misfolded proteins upon mild heat stress to protect them from degradation^21,22^. Consistently, we confirmed the co-localization of cytoplasmic Dri1 aggregates with Hsp16, a known component of PACs^21^ (Fig. S2). On the other hand, unlike Dri1, Rhs1 aggregation was not observed even at 39°C both in the wild-type and *dri1Δ* strains (Fig. 3D). Thus, it appears that, although Dri1 and Rhs1 function as a complex, only Dri1 is assembled into PACs independently of Rhs1 upon heat stress.

### The Dri1-Rhs1 complex interacts with Ccr4-Not, a eukaryotic protein complex regulating mRNA metabolism

Aiming to identify proteins physically interacting with the Dri1-Rhs1 complex, its immunoprecipitation was performed using a strain that expresses Dri1 and Rhs1 from their chromosomal loci as fusions with the *myc* and FLAG epitope tags, respectively. A tandem affinity purification of the Dri1-Rhs1 complex, followed by mass spectrometry, identified the components of the Ccr4-Not complex, Not1, Not3, and Mot2^23^ (see *Introduction*), as well as Sum3, a member of the DEAD box RNA helicase family^24^ (Fig. 4A and B). Indeed, immunoprecipitation of Dri1-myc or Rhs1-FLAG copurified GFP-tagged Not1 and Not3 (Fig. 4C-F), confirming the physical interaction between the Dri1-Rhs1 complex and Ccr4-Not. A previous mass-spectrometry study also reported Dri1 and Rhs1 as co-purified proteins with Mmi1, which physically associates with Ccr4-Not^16^.

**Fig. 4.**
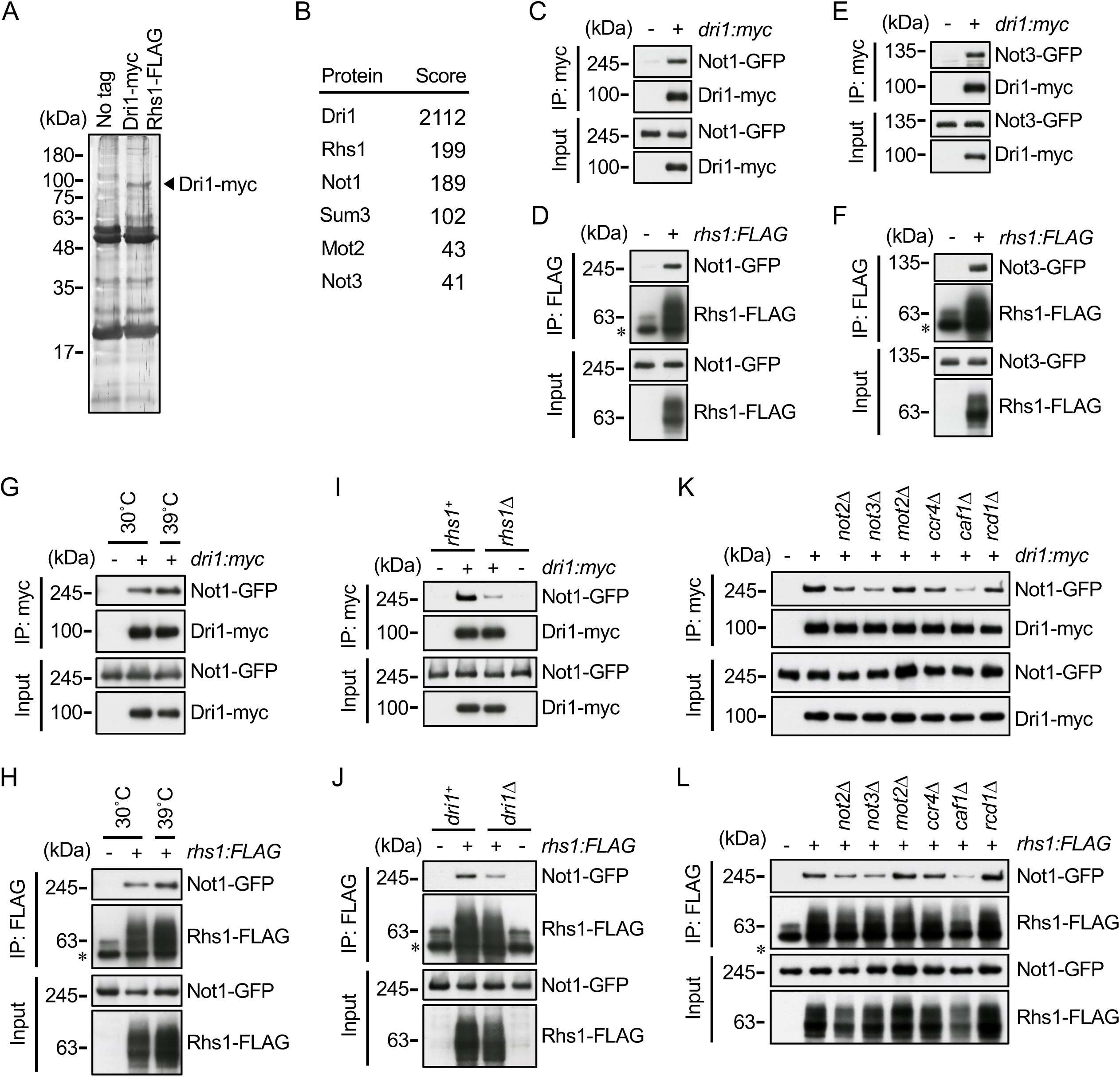
Ccr4-Not physically interacts with the Dri1-Rhs1 complex. (A) Affinity purification of the Dri1-Rhs1 complex from fission yeast. The complex was purified from the cell lysate of the *dri1:myc rhs1:FLAG* strain by two successive immunoprecipitation using anti-FLAG and anti-myc antibodies. The recovered samples were separated by SDS-PAGE followed by silver staining. A wild-type strain was used as a negative control (No tag). (B) Mass spectrometric analyses of proteins co-purified with the Dri1-Rhs1 complex shown in (A). (C-F) Physical interaction between the Dri1-Rhs1 complex and either Not1 or Not3. *dri1:myc* (C and E) or *rhs1:FLAG* (D and F) cells expressing the GFP-tagged Not1 (C and D) or Not3 (E and F) protein were cultured in YES medium at 30°C. Their cell lysate was subjected to immunoprecipitation using the anti-myc (IP: myc) or anti-FLAG antibodies (IP: FLAG), and co-purified Not1-GFP and Not3-GFP were analyzed by immunoblotting. (G and H) Interaction between the Dri1-Rhs1 complex and Not1 at high temperatures. The *dri1:myc* (G) or *rhs1:FLAG* (H) strain that expresses Not1-GFP was cultured in YES medium at 30°C and shifted to 39°C. After 4-hour incubation at 39°C, cell lysate was recovered and subjected to immunoprecipitation as in (C) and (D). (I and J) The formation of the Dri1-Rhs1 complex is required for the integrity of its interaction with Ccr4-Not. Physical interaction between Not1-GFP and Dri1-myc (I) or Rhs1-FLAG (J) in the *rhs1Δ* (I) or *dri1Δ* (J) background was analyzed as in (C) and (D). (K and L) Physical interaction between the Dri1-Rhs1 complex and Not1 in the absence of the Ccr4-Not subunits. The indicated null mutants of the Ccr4-Not subunits that express Not1-GFP and either Dri1-myc (K) or Rhs1-FLAG (L) were cultured in YES medium at 30°C, and their cell lysate was subjected to immunoprecipitation as in (C) and (D).

Further immunoprecipitation experiments found that the association between the Dri1-Rhs1 complex and Ccr4-Not is not significantly affected by the temperature shift from 30°C to 39°C (Fig. 4G and H). On the other hand, the interaction between Dri1 and Not1 was notably compromised by the *rhs1Δ* mutation (Fig. 4I). Similarly, the Rhs1-Not1 interaction was reduced in the *dri1*Δ background (Fig. 4J), indicating that the formation of the Dri1-Rhs1 complex promotes its interaction with Ccr4-Not. The association of the Dri1-Rhs1 and Ccr4-Not complexes was also examined in a series of mutants lacking individual subunits of the Ccr4-Not complex. Among the null mutants tested, *not2Δ*, *not3Δ*, and *caf1Δ* impaired the interaction of Not1-GFP with Dri1-myc and Rhs1-FLAG (Fig. 4K and L). Thus, the Not2, Not3, and Caf1 subunits appear to play crucial roles in the interaction between Ccr4-Not and the Dri1-Rhs1 complex.

### The Ccr4-Not subunits contribute to suppressing high-temperature cell growth

Because of its interaction with the Dri1-Rhs1 complex, which suppresses cellular heat resistance, Ccr4-Not was further characterized for its physiological role under high-temperature conditions. The *mot2Δ*, *caf1Δ, ccr4Δ*, and *rcd1Δ* mutants showed compromised growth even at temperatures that allow wild-type cells to grow (Fig. 5A-D). On the other hand, the *not2Δ* mutant strain exhibited better growth than the wild-type strain at 38°C (Fig. 5E). In addition, the *not3Δ* mutation significantly improved the cellular heat resistance (Fig. 5F), implying that, like the Dri1-Rhs1 complex, Ccr4-Not contributes to restricting high-temperature growth of fission yeast. To examine the functional relationships between the Dri1-Rhs1 complex and Not3, double mutants of these proteins were constructed for epistasis analyses. The high-temperature growth of the *dri1Δ not3Δ* and *rhs1Δ not3Δ* double mutants was no better than that of the *dri1Δ* and *rhs1Δ* single mutants (Fig. 5F); therefore, the heat resistance conferred by the *not3Δ* mutation is not additive to that by the *dri1Δ* and *rhs1Δ* mutations. The growth of the *dri1Δ not3Δ* and *rhs1Δ not3Δ* double mutants was slightly slower than that of the *dri1Δ* and *rhs1Δ* single mutants at any temperature, an observation attributable to the modest growth retardation caused by the *not3Δ* mutation (Fig. 5G).

**Fig. 5.**
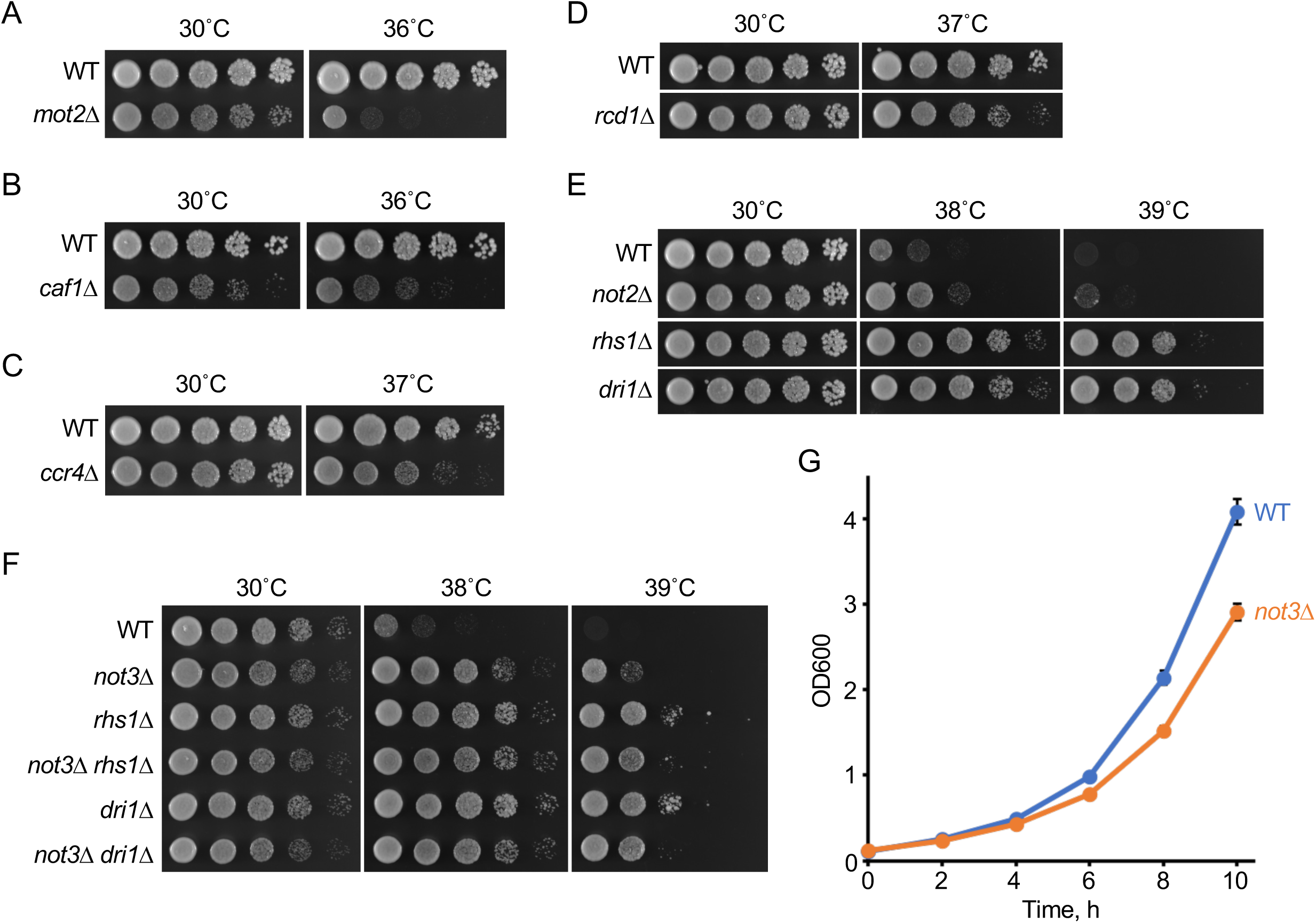
Ccr4-Not suppresses cell growth at high temperatures. (A-F) Cellular growth of the mutants lacking the subunits of the Ccr4-Not complex at high temperatures. The indicated mutant strains lacking the components of Ccr4-Not were grown in YES medium at 30°C, and their growth was tested at indicated temperatures by spotting serial culture dilutions on YES agar plates. (G) The *not3Δ* mutation impaired cell proliferation at 30°C. Wild-type and *not3Δ* cells were grown in YES liquid medium at 30°C. At 0 hour, the initial OD was adjusted to OD_600_ = 0.115 ± 0.005, and the cell growth at 30 C° was monitored by measuring OD600 at indicated time points. Data are presented as means ± s.d. from three independent experiments.

Together, these results suggest that the Dri1-Rhs1 and Ccr4-Not complexes may cooperate in suppressing cell growth at high temperatures. Among the Ccr4-Not subunits, Not2 and Not3 appear to be particularly important for this cellular function, as their absence compromises the interaction between the two complexes (Fig. 4 K and L) and improves cellular heat resistance (Fig. 5E and F).

### Dri1-Rhs1 and Ccr4-Not suppress the expression of ribosome biogenesis genes

Ccr4-Not controls gene expression by regulating various steps of mRNA metabolism, from synthesis in the nucleus to degradation in the cytoplasm^14,15^. In addition, Dri1 contains two types of RNA-binding domains, RRM and RanBP2-type zinc finger motifs. Therefore, we hypothesized that the Dri1-Rhs1 complex suppresses cellular growth at high temperatures by modulating gene expression together with Ccr4-Not. To test the possibility, we identified genes whose expression is regulated by the Dri1-Rhs1 complex. Gene expression profiling analysis by RNA-sequencing revealed that 101 and 67 genes are up-regulated, and 45 and 15 genes are down-regulated in the *dri1Δ* and *rhs1Δ* mutants, respectively (Fig. 6A, Table S1 and S2). As Dri1 and Rhs1 function as a complex in restricting cellular heat resistance, we focused on the 59 up-regulated and 11 down-regulated genes shared between the *dri1Δ* and *rhs1Δ* mutants and conducted gene ontology (GO) analysis for biological processes (Fig. 6A). While no significant enrichment of GO terms was observed in the down-regulated genes, the most up-regulated genes in the *dri1Δ* and *rhs1Δ* mutants were classified as ribosome biogenesis (Ribi) genes, which encode proteins that assist in rRNA maturation and ribosome assembly (Fig. S3). The volcano and box plots in Figures 6B-D further illustrate the significant up-regulation of Ribi genes but not ribosomal protein (RP) genes encoding ribosomal subunits. We performed RT-qPCR analysis and confirmed the increased expression of the Ribi genes *imp4^+^*, *mrd1^+^*, *cdh1^+^*, and *icp5^+^* in the *dri1Δ* and *rhs1Δ* mutants at both 30°C and 39°C (Fig. 6E). Moreover, a similar up-regulation of the Ribi genes was observed in the *not3Δ* mutant (Fig. 6E). Together, these results suggest that the Dri1-Rhs1 complex negatively regulates Ribi gene expression in conjunction with Ccr4-Not. As the physical interaction between the Dri1-Rhs1 complex and Ccr4-Not was also detected (Fig. 4), the Dri1-Rhs1 complex may recruit Ccr4-Not to Ribi gene mRNAs for their degradation (see *Discussion*).

**Fig. 6.**
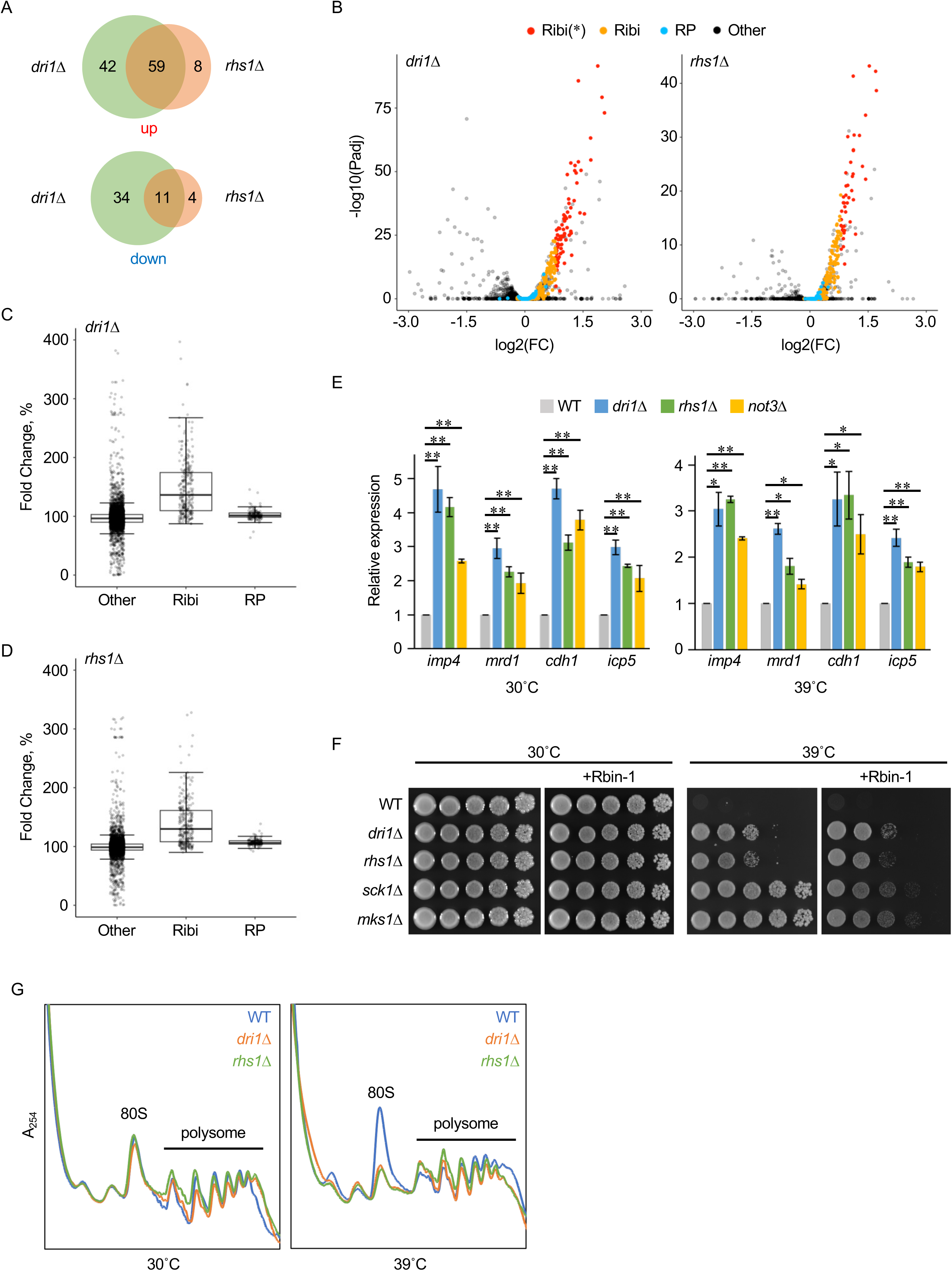
The Dri1-Rhs1 complex negatively regulates the expression of Ribi genes. (A) Venn diagram depicting the overlap of the up- and down-regulated genes between the *dri1Δ* and *rhs1Δ* mutants. (B) Volcano plots representing differentially expressed genes in the *dri1Δ* (left) and *rhs1Δ* (right) mutants compared to wild-type cells. Ribi genes and RP genes are shown in orange and blue, respectively. Ribi genes with significant difference (|log2 fold change| > 0.8; P adjusted value < 0.05) are separately illustrated in red. (C and D) Box plots illustrating the fold change of the Ribi and RP genes in the *dri1Δ* (C) and *rhs1Δ* (D) mutants. (E) Relative expression levels of representative Ribi genes (*imp4*^+^, *mrd1*^+^, *cdh5*^+^ and *icp5*^+^) in the indicated strains were determined by RT-PCR. Data represent the mean ± SD (n = 3 (30°C) and 2 (39°C)). \**P* < 0.05; \*\**P* < 0.01, compared to the wild-type control using Student’s t-test. (F) High-temperature growth of fission yeast in the presence of a ribosome biogenesis inhibitor. The indicated mutant strains were grown in YES medium at 30°C, and their growth in the presence of Rbin-1 (1 µM) was tested at indicated temperatures by spotting serial culture dilutions on YES agar plates. (G) Polysome profiles of fission yeast cells under high temperature conditions. The indicated cells were grown in EMM liquid medium at 30°C and sifted to 39°C. After 24-hour incubation at 39°C, cells were harvested and subjected to polysome analysis.

Next, we examined whether the augmented expression of Ribi genes in the *dri1Δ* and *rhs1Δ* mutants indeed contributes to their heat-resistant growth, using the ribosome biogenesis inhibitor Rbin-1^25^. The growth of fission yeast cells at 30°C was not significantly affected by Rbin-1 at the concentration tested (Fig. 6F). On the other hand, the compound impaired the heat resistance conferred by the deletion of the *sck1^+^* and *mks1^+^* genes, whose products restrict high-temperature growth independently of Dri1 and Rhs1^11^. Thus, ribosome biogenesis appears to be a critical cellular process for growth at elevated temperatures. Interestingly, the growth of *dri1Δ* and *rhs1Δ* cells at high temperatures was only marginally affected by Rbin-1 (Fig. 6F). The limited sensitivity of these mutants to Rbin-1 may be due to their induced expression of Ribi genes, which is likely to counteract the action of the drug.

Having found that the Dri1-Rhs1 and Ccr4-Not complexes modulate Ribi gene expression, we next performed polysome analysis to compare the translational process in the wild-type, *dri1Δ*, and *rhs1Δ* strains. The polysome profile in wild-type cells was comparable to that in the *dri1Δ* and *rhs1Δ* mutants at 30°C (Fig. 6G). After a temperature shift to 39°C, the 80S monosome peak was significantly elevated in wild-type cells. Although the polysome peaks were not notably affected under the stress condition, the accumulation of monosomes may represent compromised translation. Interestingly, such an increase in the 80S monosome was not detected in the *dri1Δ* and *rhs1Δ* mutants exposed to 39°C (Fig. 6G). The elevated expression of Ribi genes in these mutants may complement the translational defect detectable as monosome accumulation, thereby enabling fission yeast cells to proliferate at high temperatures.

## Discussion

Recent climate warming has increased the risk of living organisms being exposed to high temperatures that impact their growth and survival^26^. Our study, which aimed to discover the molecular mechanisms that determine cellular heat resistance, previously found that a fission yeast strain lacking either Dri1 or Rhs1 can proliferate at high temperatures inhibitory to the growth of wild-type cells^11^. In this study, we have delved into the cellular functions of these proteins to understand how they regulate the upper thermal limit of cell growth in fission yeast.

As shown in Fig. 2, the formation of the Dri1-Rhs1 complex is essential for restricting cell growth at high temperatures. It was previously reported that Dri1 physically interacts with Dpb4, a subunit of DNA polymerase epsilon, and regulates heterochromatin assembly in fission yeast^12^. In our fluorescence microscopy analysis (Fig. 3), a single, dot-like signal of Dri1 was observed within the nucleus, which may represent a fraction of Dri1 involved in heterochromatin assembly. On the other hand, most of the Dri1 protein appears diffused throughout the cytoplasm, as observed also with Rhs1. Thus, Dri1 is likely to regulate the formation of heterochromatin independently of Rhs1, and this function does not appear to contribute to the suppression of the high-temperature growth of *S. pombe*. A previous study showed that Dri1 is exported from the nucleus by the mRNA export pathway^13^. In the *rhs1Δ* mutant, Dri1 is detectable in the nucleoplasm (Fig. 3A), implying that Dri1 bound to Rhs1 is preferentially retained in the cytoplasm. While Dri1 forms a complex with Rhs1, only Dri1 is incorporated into PACs, cytoplasmic aggregates observed upon mild heat stress^21,22^. Thus, the Dri1-Rhs1 association may not be very stable, allowing Dri1 to shuttle into the nucleus and function with another partner protein, Dpb4. The significance of such dual functionality of Dri1 remains unknown.

The affinity purification of the Dri1-Rhs1 complex from fission yeast cell lysate led to the identification of Ccr4-Not as its interacting factor (Fig. 4). In addition, our gene expression analysis revealed that Dri1-Rhs1 as well as Ccr4-Not negatively regulates the expression of Ribi genes (Fig. 5). Ccr4-Not is known to control gene expression by regulating mRNA metabolism at various steps; it is involved in mRNA decay by degrading the poly(A) tail at the 3’ end of mRNA in the cytoplasm, while also regulating transcription of mRNA and its export in the nucleus^15,27^. As the Dri1-Rhs1 complex was localized in the cytoplasm (Fig. 3), we speculate that this complex downregulates the mRNA levels of Ribi genes by facilitating Ccr4-Not-dependent deadenylation. The recruitment of Ccr4-Not to particular mRNA targets for degradation is thought to be mediated by RNA-binding proteins that recognize specific sequence elements, predominantly in the untranslated regions^28,29^. For instance, human tristetraprolin (TTP), an RNA-binding protein that regulates the inflammatory response, binds to AU-rich elements in target mRNAs and recruits the Ccr4-Not complex for mRNA turnover^30,31^. Dri1 contains the RNA-binding domains, and thus, the Dri1-Rhs1 complex may mediate the binding of Ccr4-Not to the mRNAs of Ribi genes. Furthermore, the physical interaction between the Dri1-Rhs1 complex and Ccr4-Not was compromised by the *not3Δ* mutation (Fig. 4K and L), suggesting that Not3 contributes to the physical interaction of Ccr4-Not with the Dri1-Rhs1 complex. A previous study revealed that, in *Drosophila*, the RNA-binding protein Bicaudal-C (Bic-C), which is required for embryo patterning, directly binds to Not3/5, an ortholog of the fission yeast Not3, and recruits the Ccr4-Not complex to the *Bic-C* mRNA, promoting its deadenylation^32^. Therefore, the molecular function of Not3 as a module mediating the association of Ccr4-Not with RNA-binding proteins may be evolutionarily conserved across eukaryotes.

Our polysome analysis found the accumulation of 80S monosome at high temperatures in wild-type cells (Fig. 6G). On the other hand, the polysome peaks were not significantly affected under such conditions. These results indicate that protein translation is partially compromised, yet it still occurs at high temperatures, where fission yeast cells fail to proliferate. When temperature increases, cells reprogram gene expression to induce proteins required for enhanced thermotolerance, including HSPs^1,2^. It is therefore likely that proteins induced under heat stress conditions are preferentially synthesized, whereas translation of those essential for cell growth is compromised, resulting in growth defects at high temperatures. The accumulation of 80S monosome observed under high-temperature conditions is suppressed in the *dri1Δ* and *rhs1Δ* mutants (Fig. 6G). The augmented expression of Ribi genes in these mutants may suppress the translational defects of proteins required for growth at high temperatures.

We have demonstrated that the Dri1-Rhs1 complex negatively regulates Ribi gene expression in *S. pombe*. While orthologs of Rhs1 are found only in fission yeast species, Dri1 is highly conserved among eukaryotes from yeast to humans. Interestingly, the expression of budding yeast *NRP1*, a gene encoding Dri1 ortholog, is co-regulated with genes involved in the ribosome biosynthesis pathway^33^. Moreover, a previous study reported that the RanBP2-type zinc finger of TEX13A, a human ortholog of Dri1, can bind single-stranded RNA containing a GGU motif^34^. It is of great interest to understand whether those Dri1 orthologs regulate the expression of Ribi genes, as observed in fission yeast. Another intriguing question is whether the thermosensitivity of the translation process is also a determinant of the upper-temperature limit for cell growth in other eukaryotic species. Further studies in diverse eukaryotes, including functional characterization of Dri1 orthologs, will provide a better understanding of the relationship between translation and cellular thermosensitivity.

## Materials and Methods

### Fission yeast strains and general methods

*S. pombe* strains used in this study are listed in Table S3. Growth media and basic techniques for *S. pombe* have been described previously^35,36^. For the strain constructions, the PCR-based method was applied as previously reported^37,38^.

### Yeast two-hybrid assay

Yeast two-hybrid assays were performed as described previously^39,40^. Briefly, Y2HGold budding yeast strain (Clontech Laboratories) was used as a host, and interaction was determined by adenine and histidine auxotrophic markers.

### *S. pombe* growth assay

Fission yeast cells were grown in YES liquid medium, and the cultures were adjusted to the cell concentration equivalent to an optical density at 600 nm (OD_600_) of 1.0. Serial dilutions of the adjusted cultures were spotted onto agar solid media. Images were captured by the LAS-4000 system (FUJIFILM, Japan). For the growth curve assay, cells were grown in YES liquid media at 30°C. Overnight cultures were adjusted to initial OD_600_ = 0.115 ± 0.005 in fresh media, and cell density was measured at indicated time intervals.

### Immunoblotting

Crude cell lysates were prepared using trichloroacetic acid (TCA) as described previously^41^. Proteins were separated by SDS-PAGE, transferred to nitrocellulose membrane, and probed with primary antibodies as follows; anti-phospho-p70 S6K (1:5000; Cat. no. 9206, Cell Signaling Technology) for phosoho-Psk1 (Thr-415) detection, anti-DYKDDDDK (1:5000; FUJIFILM Wako) for the FLAG-tagged protein detection, anti*-myc* (1:5000; 9E10, Covance), anti-HA (1:2000; 12CA5, Roche) and anti-GFP (1:2500; Cat. no. 04404, Nacalai Tesque). Anti-rabbit IgG (H+L) HRP-conjugated (1:10000; Cat. no. W4011, Promega), anti-mouse IgG (H+L) HRP-conjugated (1:10000; Cat. no. W4021, Promega), and anti-rat IgG (H+L) HRP-conjugated (1:10000; Cat. no. 112-035-003, Jackson Immuno Research) were used as secondary antibodies.

### Immunoprecipitation and mass spectrometry

Yeast cells were disrupted in lysis buffer (20 mM HEPES-NaOH [pH 7.5], 150 mM sodium glutamate, 10% glycerol, 0.25% Tween-20, 10 mM sodium fluoride, 10 mM p-nitrophenylphosphate, 10 mM sodium pyrophosphate, 10 mM β-glycerophosphate and 0.1 mM sodium orthovanadate), containing 1 mM PMSF and protease inhibitor cocktail (P8849, Sigma-Aldrich) with glass beads using Multi-beads Shocker (Yasui Kikai). The cell lysate was recovered by centrifugation for 15 min at 17,700 × g and the total protein concentrations of cell lysates were determined by Bradford assay. For interaction between myc-tagged Dri1 and FLAG-tagged Rhs1, the recovered cell lysates were incubated with anti-FLAG M2-affinity gel (Sigma-Aldrich) for 2 h at 4°C, followed by extensive washes with lysis buffer. Resultant samples were subjected to immunoblotting.

For interaction of GFP-tagged proteins, the cell lysates were incubated with anti-DYKDDDDK (FUJIFILM Wako) or anti-myc (Covance) antibodies for 1 h at 4°C, followed by incubation with Dynabeads Protein G (Thermo Fisher Scientific) for 2 h at 4°C.

For identification of proteins interacting with the Dri1-Rhs1 complex, cell lysates were first incubated with anti-FLAG M2-affinity gel for 2 h at 4°C, followed by elution using 3X FLAG peptides (Sigma-Aldrich). The recovered samples were then further incubated with EZview^TM^ Red anti-c-Myc affinity gel for 2 h at 4°C. After extensive washes with lysis buffer, the resultant samples were resolved in a 12% Mini-PROTEAN TGX precast gel (Bio-Rad). Each lane was sliced into four pieces and digested with trypsin. Mass spectrometric analysis was performed using the LTQ-Orbitrap XL-HTC-PAL system. MS/MS spectra were analyzed by Mascot server (Matrix Science) and compared against NCBInr protein database (Taxonomy: *S. pombe*).

### Microscopy

Fluorescence microscopic analysis was performed using DeltaVision Elite Microscopy System (GE Healthcare) as described previously^40,42,43^. Briefly, cells grown exponentially in EMM liquid were stained with Hoechst 33342 (FUJIFILM Wako) for nuclear visualization and mounted on a thin layer of EMM agar. Z-axial images were taken at 0.4 μm with a 60× objective lens. Deconvolution of images was performed using DeltaVision SoftWoRx software.

### RNA extraction, quantitative RT-PCR (qPCR) and RNA-seq

Cells were grown to exponential phase at 30°C and shifted to 39°C for heat treatment experiment. After 16-hour incubation at 39°C, cells were harvested and flash-frozen using liquid nitrogen. Total RNA was purified using FastGene^TM^ Premium Kit (Nippon Genetics, Japan) according to the manufacturer’s protocol.

For RT-qPCR, the SuperScript II reverse transcriptase (Thermo Fisher Scientific) was used to synthesize cDNA. RT-qPCR was carried out using PowerUp SYBR Green Master mix (Thermo Fisher Scientific). The results were normalized against the housekeeping gene *act1^+^*. PCR primer sets used in RT-qPCR are listed in Table S4.

For RNA-seq, mRNA was enriched using oligo dT beads, and mRNA-sequencing was performed by BGI JAPAN (Kobe, Japan) using a DNBseq platform. Clean reads were mapped to reference genome using Bowtie2^44^, and gene expression level was calculated with RSEM^45^. The up-regulated and down-regulated genes were determined by DEseq2 and PossionDis algorithms^46,47^. The reference genome sequence and gene annotation information of *S. pombe* were obtained from the PomBase database (http://www.pombase.org)^48^. GO categories of the target genes were determined using GO Term Finder (https://go.princeton.edu/cgi-bin/GOTermFinder), and the volcano and box plots were generated using R studio using ggplot2 package^49^.

### Polysome analysis

Cells were grown in EMM at 30°C and shifted to 39°C for heat treatment experiment. After 24-hour incubation at 39°C, cell cultures were mixed with 0.1 mg/ml of cycloheximide and incubated 30°C for 5 min. Cells were harvested and washed with polysome lysis buffer (20 mM Tris-HCI [pH 7.5], 150 mM KCl, 10 mM MgCl_2_, 100 µg/ml of cycloheximide, 1 mM dithiothreitol, 0.2 mg/ml heparin, 1 mM PMSF, RNasin (Promega) and protease inhibitor cocktail), followed by flash-freezing with liquid nitrogen. Polysome analysis was performed using a gradient master and fractionator (107–201 M and 152–002; BioComp Instruments) as previously described^50,51^.

## Data Availability

The RNA sequence data in this article have been deposited in the DDBJ Sequence Read Archive (DRA) under accession codes DRR654225-DRR654230.

## Acknowledgments

We thank A Higuchi and the NAIST Life Science Collaboration Center (LiSCo) for technical assistance. We are also grateful to T Toda and M Yukawa for sharing unpublished data and strains.

## Competing Interests

The authors declare no competing interests.

## Author Contributions

Y.M., and K.S., designed research; Y.A., A.Y., T.N., Y.F., Y.N., F.M., S.I., and Y.M., performed research; Y.A., A.Y., T.N., Y.N., F.M., S.I., and Y.M., analyzed data; and K.S., and Y.M., wrote the paper.

## Funding

This study was supported by research grants to Y.M. from the Sumitomo Foundation and the Noda Institute for Scientific Research, and to Y.N. from the Tojuro Iijima Foundation for Food Science and Technology. This work was also supported by the Takeda Science Foundation to K.S. and Y.M., the Ohsumi Frontier Science Foundation to S.I. and K.S., and the Institute for Fermentation, Osaka (IFO) to S.I. and Y.M, and the Japan Society for the Promotion of Science (JSPS) KAKENHI grants (19K06564, 22K06145 and 25K09615 to Y.M., 20K06485 and 25K09040 to Y.N., and 19H03224 to K.S.).

## Notes

### Competing Interest Statement

The authors have declared no competing interest.

