## Supplementary Information for "Thermosensitivity of cellular translation restricts the growth of fission yeast at high temperatures"

| No. | AD | DBD |
|---|---|---|
| 1 | Rhs1 | Dri1 |
| 2 | Dri1 | Rhs1 |
| 3 | – | Dri1 |
| 4 | Dri1 | – |
| 5 | – | Rhs1 |
| 6 | Rhs1 | – |

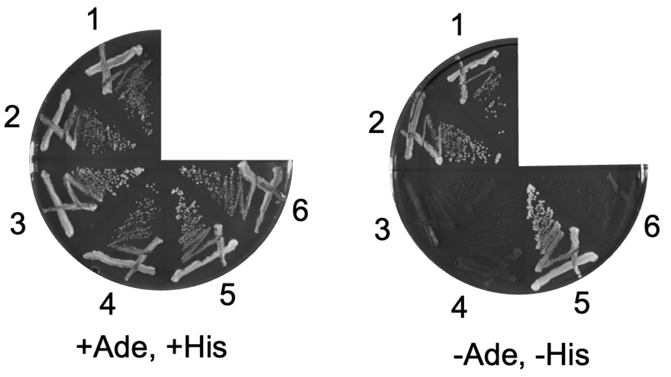


**Supplementary Figure S1. Characterization of the interaction between Dri1 and Rhs1 by yeast two-hybrid assays (related to Figure 1).**

Dri1 and Rhs1 fused to either Gal4 DNA-binding (DBD) or activation (AD) domains were expressed in the budding yeast Y2HGold (left), and their interaction was judged by adenine and histidine auxotrophy. Cells carrying either of the empty vectors were used as negative controls.

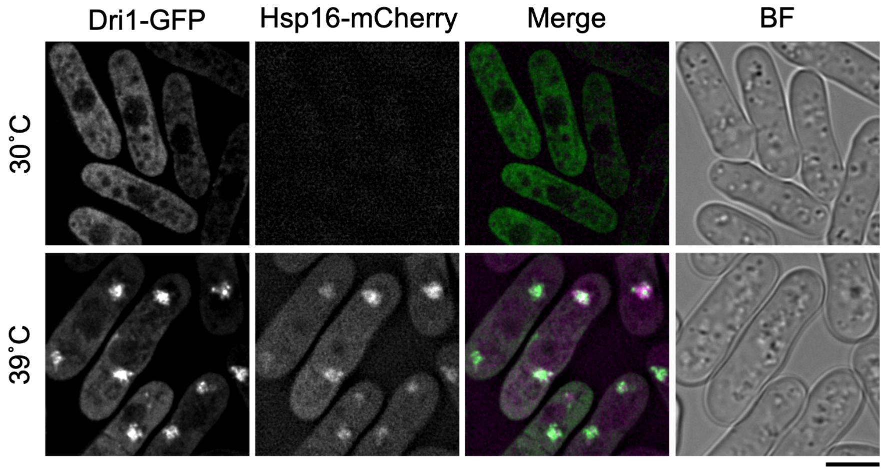


**Supplementary Figure S2. Dri1 is assembled into PACs at 39˚C**

Fission yeast strains expressing GFP-tagged Dri1 and mCherry-tagged Hsp16 from their chromosomal locus were grown in EMM at 30˚C and shifted to 39˚C. After 4-hour incubation at 39˚C, the localization of Dri1 and Hsp16 was analyzed by fluorescence microscopy. Z-axial images were collected and mid-section images after deconvolution are shown. BF, bright-field image. Bar, 5 µm.

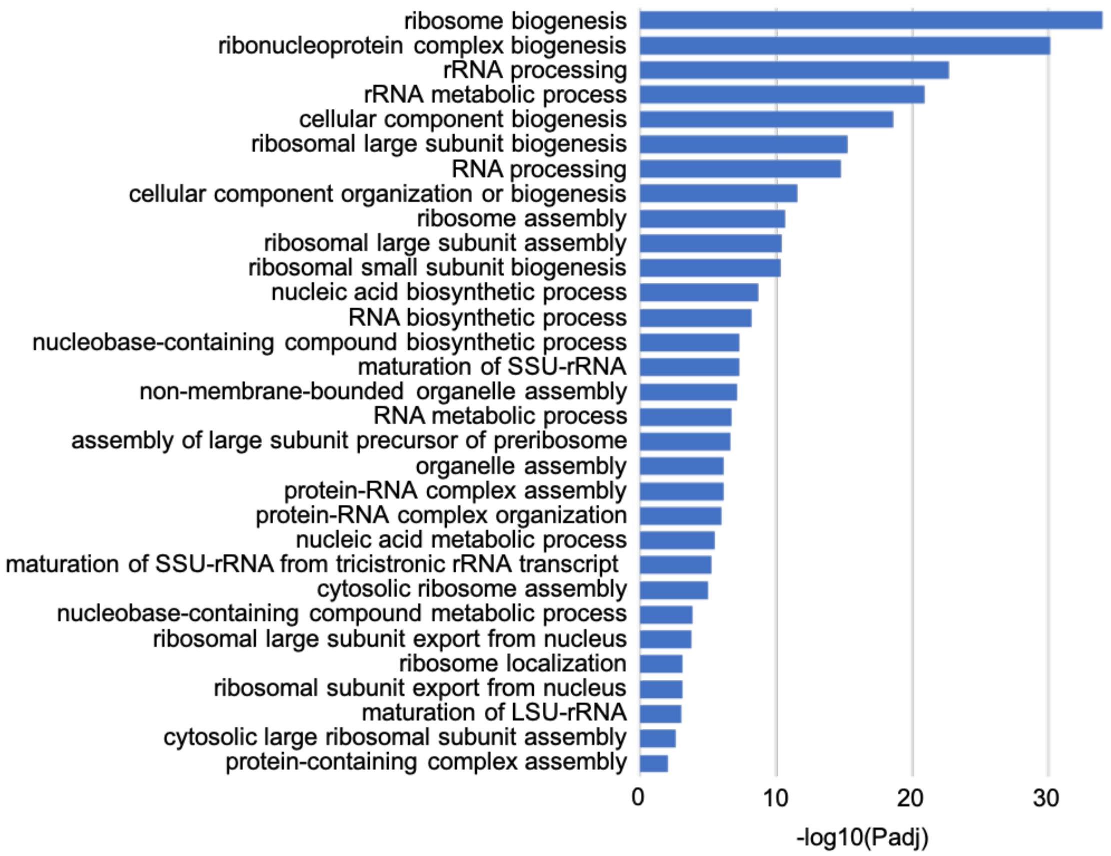


**Supplementary Figure S3. GO analysis for biological processes of the genes up-regulated in both the *dri1*Δ and *rhs1*Δ mutants**

The x-axis indicates adjusted P values in logarithmic scale.

**Supplementary Table S3.** ***S. pombe*** **strains list used in this study**

| Strain ID | Genotype | Source, Reference |
|---|---|---|
| CA15458 | *h-* | Lab stock |
| CA2 | *h+* | Lab stock |
| CA16908 | *h- dri1:13myc:hph* | Morozumi et al.,2024 |
| CA16915 | *h- rhs1:5FLAG:KanMX6:* | Morozumi et al.,2024 |
| CA16980 | *h+ dri1:13myc:hph rhs1:5FLAG:KanMX6* | Morozumi et al.,2024 |
| CA18233 | *h- dri1:13myc:hph rhs1(1-120):5FLAG:KanMX6* | This study |
| CA18234 | *h- dri1:13myc:hph rhs1(1-240):5FLAG:KanMX6* | This study |
| CA18235 | *h- dri1:13myc:hph rhs1(121-240):5FLAG:KanMX6* | This study |
| CA18236 | *h- dri1:13myc:hph rhs1(121-368):5FLAG:KanMX6* | This study |
| CA18237 | *h- dri1:13myc:hph rhs1(241-368):5FLAG:KanMX6* | This study |
| CA16856 | *h- rhs1::KanMX4* | Bioneer |
| CA18065 | *h+ rhs1(1-120):5FLAG:KanMX6* | This study |
| CA18067 | *h+ rhs1(1-240):5FLAG:KanMX6* | This study |
| CA18069 | *h+ rhs1(121-240):5FLAG:KanMX6* | This study |
| CA18071 | *h+ rhs1(121-368):5FLAG:KanMX6* | This study |
| CA18073 | *h+ rhs1(241-368):5FLAG:KanMX6* | This study |
| CA18238 | *h- dri1:13myc:hph rhs1(31-368):5FLAG:KanMX6* | This study |
| CA18239 | *h- dri1:13myc:hph rhs1(61-368):5FLAG:KanMX6* | This study |
| CA17022 | *h- dri1:mEGFP:KanMX6* | This study |
| CA17589 | *h- dri1:mEGFP:KanMX6 rhs1:5FLAG:KanMX6* | This study |
| CA18243 | *h- dri1(1-410):mEGFP:KanMX6 rhs1:5FLAG:KanMX6* | This study |
| CA17591 | *h- dri1(1-330):mEGFP:KanMX6 rhs1:5FLAG:KanMX6* | This study |
| CA17590 | *h- dri1(1-225):mEGFP:KanMX6 rhs1:5FLAG:KanMX6* | This study |
| CA20479 | *h- dri1(101-596):mEGFP:KanMX6 rhs1:5FLAG:KanMX6* | This study |
| CA20481 | *h+ dri1(218-596):mEGFP:KanMX6 rhs1:5FLAG:KanMX6* | This study |
| CA20483 | *h- dri1(323-596):mEGFP:KanMX6 rhs1:5FLAG:KanMX6* | This study |
| CA16783 | *h- dri1::KanMX4* | Bioneer |
| CA18122 | *h- dri1(1-507):EGFP:KanMX6* | This study |
| CA18121 | *h- dri1(1-402):EGFP:KanMX6* | This study |
| CA17417 | *h- dri1(1-322):EGFP:KanMX6* | This study |
| CA17415 | *h- dri1(1-217):EGFP:KanMX6* | This study |
| CA18960 | *h- dri1(101-596):EGFP:KanMX6* | This study |
| CA18962 | *h- dri1(218-596):EGFP:KanMX6* | This study |
| CA18963 | *h- dri1(323-596):EGFP:KanMX6* | This study |
| CA18969 | *h- dri1(218-507):EGFP:KanMX6* | This study |
| CA18967 | *h- dri1(218-402):EGFP:KanMX6* | This study |
| CA18965 | *h- dri1(218-328):EGFP:KanMX6* | This study |

| | | |
|---|---|---|
| CA18306 | *h- dri1:13myc:hph rhs1(FQF34,37,38AAA):5FLAG:KanMX6* | This study |
| CA18307 | *h- dri1:13myc:hph rhs1(FQ34,37AA):5FLAG:KanMX6* | This study |
| CA18308 | *h+ dri1:13myc:hph rhs1(FF34,38AA):5FLAG:KanMX6* | This study |
| CA18677 | *h+ dri1:13myc:hph rhs1(F34A):5FLAG:KanMX6* | This study |
| CA18679 | *h+ dri1:13myc:hph rhs1(Q37A):5FLAG:KanMX6* | This study |
| CA18681 | *h- dri1:13myc:hph rhs1(F38A):5FLAG:KanMX6* | This study |
| CA18202 | *h- rhs1(FQF34,37,38AAA):5FLAG:KanMX6* | This study |
| CA18204 | *h- rhs1(FQ34,37AA):5FLAG:KanMX6* | This study |
| CA18206 | *h- rhs1(FF34,38AA):5FLAG:KanMX6* | This study |
| CA18622 | *h- rhs1(F34A):5FLAG:KanMX6* | This study |
| CA18624 | *h- rhs1(Q37A):5FLAG:KanMX6* | This study |
| CA18626 | *h- rhs1(F38A):5FLAG:KanMX6* | This study |
| CA17162 | *h- dri1:mEGFP:KanMX6 rhs1::KanMX4* | This study |
| CA18618 | *h- rhs1:GBP:mCherry:KanMX6* | This study |
| CA18747 | *h- dri1:mEGFP:KanMX6 rhs1:GBP:mCherry:KanMX6* | This study |
| CA18668 | *h- rhs1(F34A):GBP:mCherry:KanMX6* | This study |
| CA18751 | *h- dri1:mEGFP:KanMX6 rhs1(F34A):GBP:mCherry:KanMX6* | This study |
| CA18672 | *h- rhs1(Q37A):GBP:mCherry:KanMX6* | This study |
| CA18754 | *h+ dri1:mEGFP:KanMX6 rhs1(F34A):GBP:mCherry:KanMX6* | This study |
| CA17442 | *h- dri1:mEGFP:KanMX6 hsp16:mCherry:KanMX6* | This study |
| CA17024 | *h- rhs1:mEGFP:KanMX6* | This study |
| CA17030 | *h- dri1::KanMX4 rhs1:mEGFP:KanMX6* | This study |
| CA18170 | *h- not1:mEGFP:KanMX6* | This study |
| CA18220 | *h- dri1:13myc:hph not1:mEGFP:KanMX6* | This study |
| CA18216 | *h- rhs1:5FLAG:KanMX6 not1:mEGFP:KanMX6* | This study |
| CA18884 | *h- not3:mEGFP:KanMX6* | This study |
| CA18902 | *h- dri1:13myc:hph not3:mEGFP:KanMX6* | This study |
| CA18900 | *h- rhs1:5FLAG:KanMX6 not3:mEGFP:KanMX6* | This study |
| CA19104 | *h- dri1:13myc:hph not1:mEGFP:KanMX6 rhs1::KanMX4* | This study |
| CA19106 | *h- rhs1:5FLAG:KanMX6 not1:mEGFP:KanMX6 dri1::KanMX4* | This study |
| CA19311 | *h- dri1:13myc:hph not1:mEGFP:KanMX6 not2::hph* | This study |
| CA19273 | *h- dri1:13myc:hph not1:mEGFP:KanMX6 not3::hph* | This study |
| CA19562 | *h- dri1:13myc:hph not1:mEGFP:KanMX6 mot2::hph* | This study |
| CA19713 | *h- dri1:13myc:hph not1:mEGFP:KanMX6 ccr4::KanMX4* | This study |
| CA19717 | *h- dri1:13myc:hph not1:mEGFP:KanMX6 caf1::hph* | This study |
| CA19566 | *h- dri1:13myc:hph not1:mEGFP:KanMX6 rcd1::hph* | This study |
| CA19313 | *h- rhs1:5FLAG:KanMX6 not1:mEGFP:KanMX6 not2::hph* | This study |
| CA19275 | *h- rhs1:5FLAG:KanMX6 not1:mEGFP:KanMX6 not3::hph* | This study |
| CA19561 | *h+ rhs1:5FLAG:KanMX6 not1:mEGFP:KanMX6 mot2::hph* | This study |
| CA19711 | *h- rhs1:5FLAG:KanMX6 not1:mEGFP:KanMX6 ccr4::KanMX4* | This study |
| CA19715 | *h- rhs1:5FLAG:KanMX6 not1:mEGFP:KanMX6 caf1::hph* | This study |

| | | |
|---|---|---|
| CA19564 | *h- rhs1:5FLAG:KanMX6 not1:mEGFP:KanMX6 rcd1::hph* | This study |
| CA18325 | *h- mot2::hph* | This study |
| CA18890 | *h- caf1::hph* | This study |
| CA18377 | *h- ccr4::KanMX4* | Bioneer |
| CA18892 | *h- rcd1::hph* | This study |
| CA18888 | *h- not2::hph* | This study |
| CA18886 | *h- not3::hph* | This study |
| CA18910 | *h- dr1i::KanMX4 not3::hph* | This study |
| CA18904 | *h- rhs1::KanMX4 not3::hph* | This study |
| CA7595 | *h- sck1::KanMX4* | Bioneer |
| CA16251 | *h- mks1::KanMX4* | Bioneer |

**Supplementary Table S4. Primer list used in RT-qPCR**

| Primer name | sequence |
|---|---|
| act1_F | 5'-AAGTACCCCATTGAGCACGG-3' |
| act1_R | 5'-CAGTCAACAAGCAAGGGTGC-3' |
| imp4_F | 5'-TTCCGTCACCACATTTACGC-3' |
| imp4_R | 5'-TGTGACGCTGGTAAGGCTTC-3' |
| mrd1_F | 5'-ACTCGGTGCTTATGGGCAAC-3' |
| mrd1_R | 5'-AGAGCTCTCATGGCATTGGC-3' |
| cdh1_F | 5'-CGAACGCATGGTTGAGGAAG-3' |
| cdh1_R | 5'-GAACGCCAACCTGATGGAAG-3' |
| icp5_F | 5'-GCTTGATCGCAAATTCCATG-3' |
| icp5_R | 5'-CTGGCCAATTTGAACAGGTG-3' |
